# Integrative analysis of pharmacogenomics in major cancer cell line databases using CellMinerCDB

**DOI:** 10.1101/292904

**Authors:** Vinodh N. Rajapakse, Augustin Luna, Mihoko Yamade, Lisa Loman, Sudhir Varma, Margot Sunshine, Francesco Iorio, Fabricio G. Sousa, Fathi Elloumi, Mirit I. Aladjem, Anish Thomas, Chris Sander, Kurt Kohn, Cyril H. Benes, Mathew Garnett, William C. Reinhold, Yves Pommier

**Affiliations:** Developmental Therapeutics Branch, Center for Cancer Research, National Cancer Institute, NIH, Bethesda, MD 20892.; cBio Center, Dana-Farber Cancer Institute and Department of Cell Biology, Harvard Medical School, Boston, MA; Wellcome Trust Sanger Institute, Wellcome Trust Genome Campus, Hinxton, UK.; Massachusetts General Hospital Cancer Center and Department of Medicine, Harvard Medical School, Charlestown, Massachusetts, USA.; First Department of Medicine, Hamamatsu University School of Medicine, Hamamatsu, Japan.; Centro De Estudos Em Células Tronco, Terapia Celular E Genética Toxicológica, Programa De Pós-Graduação Em Farmácia, Universidade Federal De Mato Grosso Do Sul, Campo Grande, MS 79070-900, Brazil

## Abstract

As precision medicine demands molecular determinants of drug response, CellMinerCDB provides (https://discover.nci.nih.gov/cellminercdb/) a web-based portal for multiple forms of pharmacological, molecular, and genomic analyses, unifying the richest cancer cell line datasets (NCI-60, NCI-SCLC, Sanger/MGH GDSC, and Broad CCLE/CTRP). CellMinerCDB enables genomic and pharmacological data queries for identifying pharmacogenomic determinants, drug signatures, and gene regulatory networks for researchers without requiring specialized bioinformatics support. It leverages overlaps of cell lines and tested drugs to allow assessment of data reproducibility. It builds on the complementarity and strength of each dataset. A panel of 41 drugs evaluated in parallel in the NCI-60 and GDSC is reported, supporting drug reproducibility across databases, repositioning of bisacodyl and acetalax for triple negative breast cancer, and identifying novel drug response determinants and genomic signatures for topoisomerase inhibitors and schweinfurthins in development. CellMinerCDB also allowed the identification of *LIX1L* as a novel mesenchymal gene regulating cellular migration and invasiveness.

---

A critical aim of precision medicine is to match drugs with genomic determinants of response. Associating tumor sample molecular features with the response to specific drug treatments is especially challenging because of the typically encountered diversity of patient experiences, partial knowledge of the multiple molecular determinants of response and resistance downstream from the primary drug targets, and tumor heterogeneity. In this setting, the relative homogeneity of cell populations in cancer cell lines is advantageous, making them a starting point for resolving and establishing cell intrinsic drug response mechanisms. These features motivate the expanded development of cancer cell line pharmacogenomic databases.

Building on the paradigm introduced by the NCI-60 (1, 2), pharmacogenomic data portals such the Genomics of Drug Sensitivity in Cancer (GDSC) (3, 4), the Cancer Cell Line Encyclopedia (CCLE) (5), and the Cancer Therapeutics Response Portal (CTRP) (6) have expanded to span ~1000 cancer cell lines. Each database provides a readily available resource for translational research, and proposals have been advanced to further enrich them to over 10,000 cancer cell lines for better coverage of tumor type diversity (7). The CellMiner NCI-60 dataset includes drug activity data for over 21,000 compounds, together with a wide range of molecular profiling data (gene expression, mutations, copy number, methylation, and protein expression). The GDSC and CCLE collections focus on drug activity data for clinically relevant drugs over larger cell line sets, together with an array of molecular profiling data that match the NCI-60 and clinical genomic analyses. The CTRP provides independent drug activity data for nearly 500 compounds over cell lines spanning most of the CCLE and GDSC collections. The source-specific portals allow deep exploration of their associated data sets, but they do not allow immediate cross-database analyses. Yet, substantial overlaps in both cell lines and drugs have the potential to allow integrative analyses, building on the complementarity of the cancer cell line data sets. But data complexity and mundane sources of friction, such as differently named entities (cell lines, drugs), has until now made working across databases challenging, even for those with informatics training.

To unleash the power of integrative analyses within and across data sources, we have developed CellMinerCDB (https://discover.nci.nih.gov/cellminercdb/), a web application allowing immediate, interactive exploration of cancer cell line pharmacogenomic data (Figure 1). In CellMinerCDB, named entities are transparently matched across sources, allowing cell line molecular features and drug responses to be readily compared using bivariate scatter plots and correlation analyses. Multivariate models of drug response or any other genomic cell line attribute can be assessed. Analyses can be restricted to tissues of origin, with cell lines across all sources mapped to a uniform tissue type hierarchy. Gene pathway annotations allow assessment and filtering of analysis results. Moreover, CellminerCDB is built using the publicly available rcellminer R/Bioconductor package, which provides analyses and a standard data representation format (8). The latter also allows CellMinerCDB to be readily updated to include additional data.

**Figure 1:**
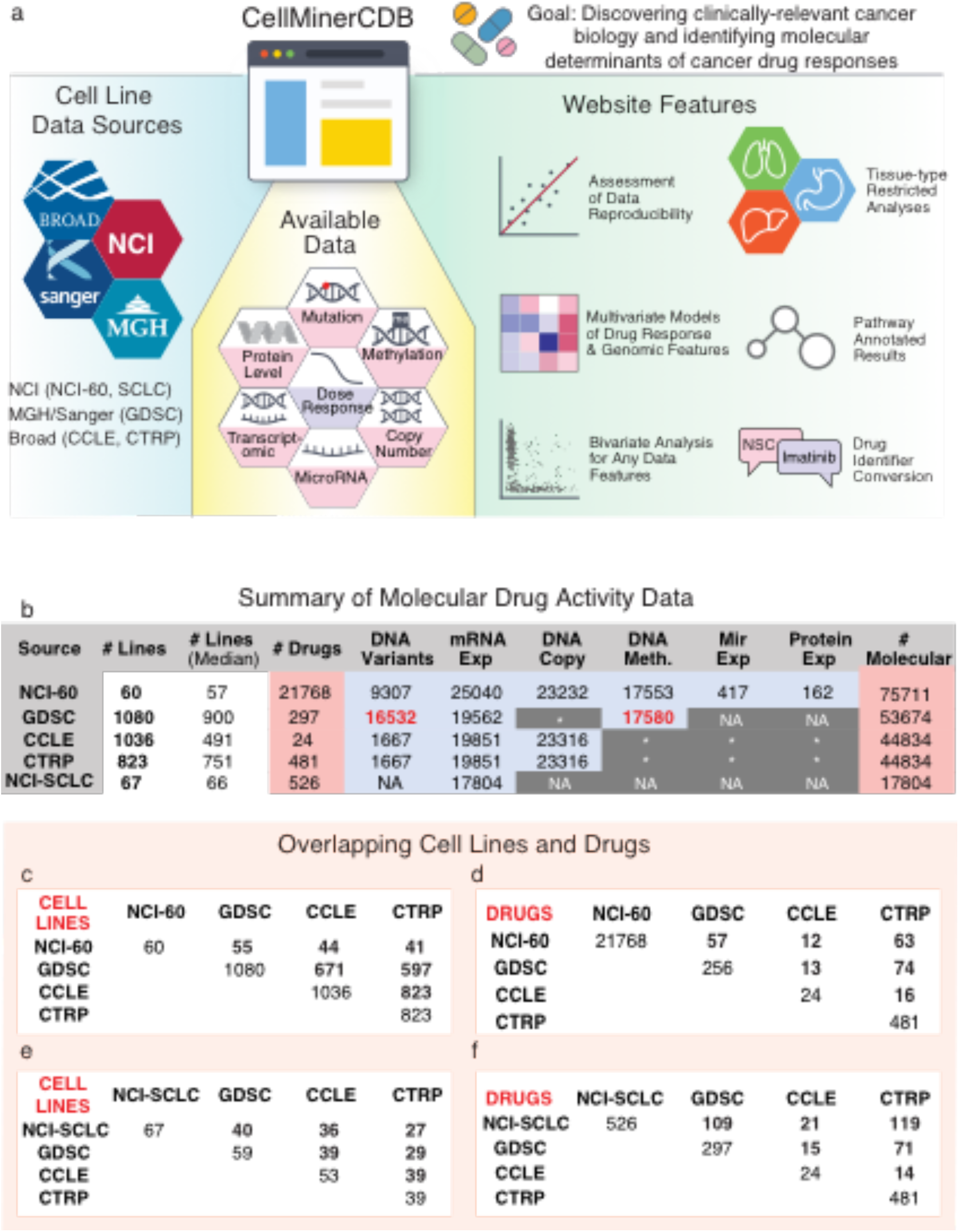
CellMinerCDB. integrates cancer cell line information from principal resources and provides powerful, user-friendly analysis tools (a). **Data source comparisons.** (b) Summary of molecular and drug activity data for the 5 data sources currently included in CellMinerCDB. For molecular data types, the numbers indicate the number of genes with a particular data type. GDSC gene-level mutation and methylation data (numbers in red) were prepared from raw data as part of the development of CellMinerCDB. Asterisks indicate molecular data under development, but not publicly available. (c - f) Cell line and drug overlaps between data sources. Protein expression was determined by RPPA (Reverse-Phase Protein Array).

In this report, we present CellMinerCDB and key features of molecular and drug data reproducibility, and complementarity across sources. We provide examples illustrating cancer biology explorations and drug response determinants. These include multivariate modeling of the response to topoisomerase inhibitors and to Schweinfurthins, a class of NCI-developed compounds derived from natural products with novel mechanism of action. CellMinerCDB also provides phenotypic signatures for cancer cell lines, including a gene expression-based measure of epithelial-to-mesenchymal (EMT) transition status. We use the latter to assess EMT stratification within specific tissues of origin, and to identify a novel EMT gene, *LIX1L*. Detailed use of CellMinerCDB is described in a video tutorial (https://youtu.be/2HicAgcyJHI).

## Results

### Data source comparisons

CellMinerCDB integrates four prominent cancer cell line data sources (CellMiner NCI-60 (2,8–10), Sanger/Massachusetts General Hospital GDSC (3), the Broad/Novartis CCLE, and the Broad CTRP) (5, 6) and a tissue-specific dataset encompassing 66 small cell lung cancer lines (NCI SCLC) (11) (Figure 1). Collectively, these sources provide drug activity and molecular profiling data for approximately 1,400 distinct cancer cell lines (Figure 1b, Supplementary Data). Each source has particular strengths. The NCI-60 is unmatched with respect to breadth of molecular profiling data, as well as the number of tested drugs. The NCI-60 data also include replicate data readily accessible via the established CellMiner data portal (2). The GDSC, CCLE, and CTRP sources feature much larger numbers of cell lines, spanning tissues of origin not included in the NCI-60. The range of tested compounds in these expanded cell line panels is narrow relative to the NCI-60, though the GDSC and CTRP focus on a wide range of clinically relevant anticancer drugs. The CTRP provides data for 170 FDA-approved or investigational anticancer drugs and 196 other compounds with mechanism of action information. The CTRP molecular data in CellMinerCDB are from the CCLE (Figure 1b).

Despite ongoing data acquisition and processing efforts, gaps exist with respect to genomic profiling data in sources other than the CellMiner NCI-60 (Figure 1b, dark grey table entries). For the GDSC gene mutation and methylation data, we took advantage of processing pipelines developed for the NCI-60 (10, 12) to compute gene-level summary data from publicly available raw data. Remaining source-specific molecular profiling data gaps can be filled within CellMinerCDB by effectively extending data provided by one source to another. This is possible because of extensive overlaps between tested cell lines and drugs (Figures 1c-f). For example, methylation data are not publicly available for the CCLE, but GDSC methylation data are available for the matching 671 cell CCLE lines and 597 CTRP lines (Figure 1c). CellMinerCDB automatically matches synonymous cell line and drug names, freeing users from a mundane impediment to work across data sources.

### Molecular data reproducibility

Integrative analyses presuppose data concordance across sources. Such analyses can be readily performed with CellMinerCDB because of the large overlaps across the cancer cell line databases. For instance, 55 of the NCI-60 lines are in GDSC and 44 are in CCLE, while 671 lines (~60%) are shared between CCLE and GDSC (Figure 1b); 40 of the 67 NCI-SCLC lines are in GDSC and 36 are in CCLE (Figure 1e); 74 drugs are in both GDSC and CTRP, and 63 drugs are in both NCI-60 and CTRP (Figure 1d).

For molecular data, we assessed data concordance by computing Pearson’s correlations between gene-specific molecular profiles over matched cell lines for all pairs of sources and comparable data types. The distributions of expression, copy number, and methylation data correlations indicate substantial concordance across sources (Figure 2a). For these analyses, gene-level transcript expression and methylation patterns with uniformly low values across matched cell lines from both compared sources were excluded (Supplementary Methods). While the median number of matched cell lines varies widely over the compared sources and molecular data types, the median correlations exceed 0.7 in all cases (Figure 2a). The striking concordance between NCI-60 and GDSC methylation data (median R = 0.97, median n = 52) may derive in part from the use of same technology platform (10) and gene-level data summarization approach for each source (Materials and Methods). Examples for specific genes are displayed in Supplementary Figure 1 demonstrating the high data reproducibility for *SLFN11* (Schlafen 11) expression in the NCI-60 vs. GDSC, *CDH1* (E-cadherin) expression in GDSC vs. CCLE, *SLFN11* methylation in the GDSC vs. NCI-60 and *CDKN2A* (p16^INK4^/p19^ARF^) copy number in NCI-60 vs. CCLE. Readers are invited to explore their own queries at https://discover.nci.nih.gov/cellminercdb/ by selecting a genomic feature for any given gene in two different datasets of their choice.

**Figure 2:**
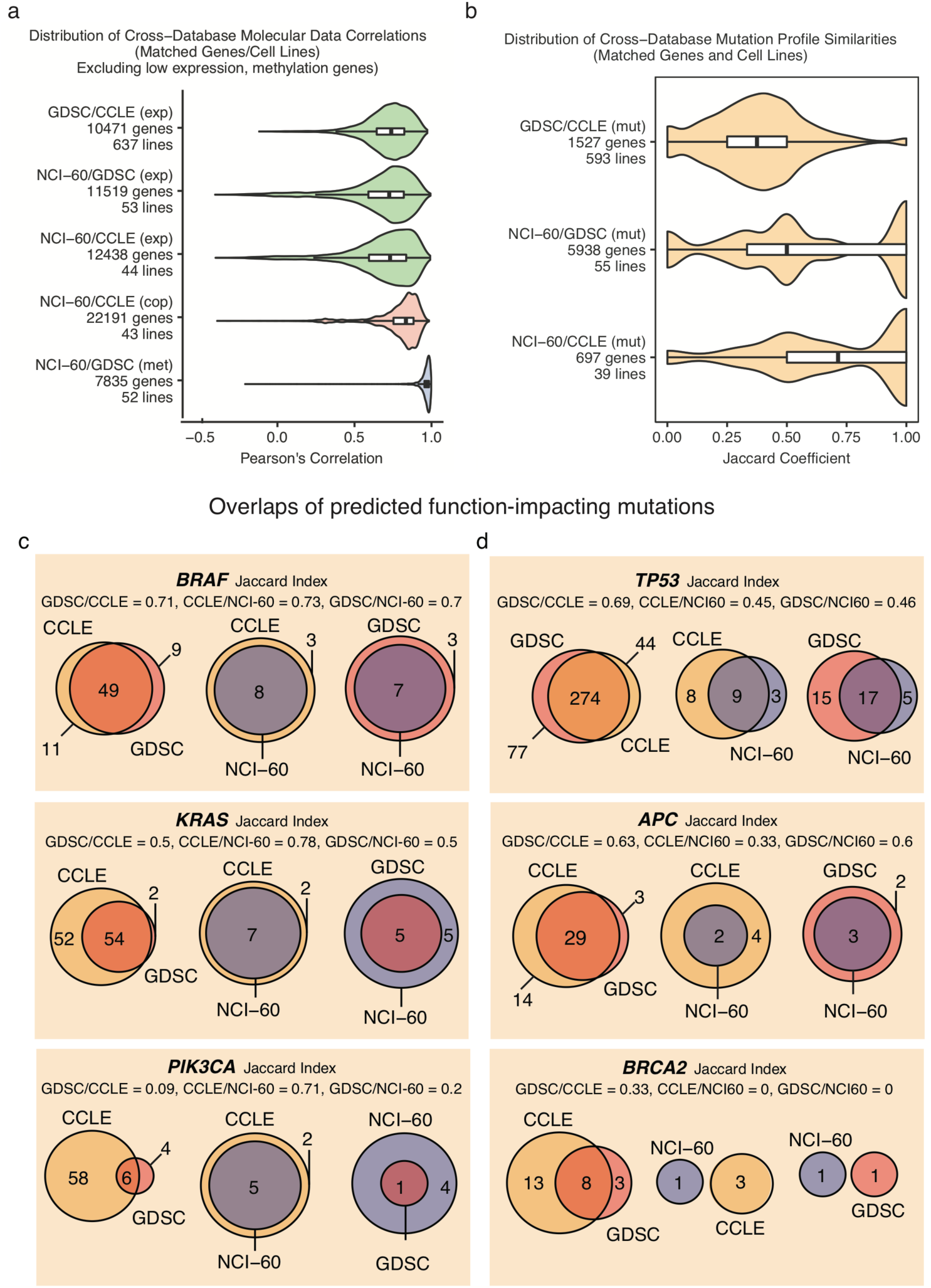
Molecular data reproducibility across sources. Comparison of the available genomic features of the cell lines shared between the CellMinerCDB data sources. Bar plots indicate the median and inter-quartile range. (a) Pearson’s correlation distributions for comparable expression (exp), DNA copy number (cop), and DNA methylation (met) data. (b) Jaccard coefficient distributions for comparable binary mutation (mut) data. The Jaccard coefficient for a pair of gene-specific mutation profiles is the ratio of the number of mutated cell lines reported by both sources to the number of mutated lines reported by either source. (c, d) Overlaps of function-impacting mutations as predicted using SIFT/PolyPhen2 for selected tumor suppressor genes and oncogenes. Matched cell line mutation data were binarized by assigning a value of one to lines with a homozygous mutation probability greater than a threshold, which was set to 0.3 for (b) and for oncogenes in (c), and to 0.7 for tumor suppressor genes in (d).

Gene-level mutation data in CellMinerCDB indicate the probability that an observed mutation is homozygous and is function impacting. These values were converted to cumulative probability values for genes with multiple deleterious mutations (12), and are available in graphical and tabular forms at https://discover.nci.nih.gov/cellminercdb/. To compare gene-specific mutation profiles across sources, we binarized the matched cell line data by assigning a value of 1 to lines with an aforementioned probability value greater than 0.3. Entirely matched mutation profiles across sources should have a Jaccard index value of 1. As such, the similarity index distributions indicate greater discordance for the mutation data (Figure 2b) than for the other types of genomic data (Figure 2a). The similarity distribution values are higher for comparisons with the NCI-60 data (NCI-60/GDSC median J = 0.5, n = 55; NCI-60/CCLE median J = 0.71, n = 39) than for the GDSC/CCLE comparison (median J = 0.38, n = 593). One caveat, however, is that the large cell line data base comparisons entail far larger numbers of matched cell lines. Indeed, the Jaccard similarity values approaching 1 with the NCI-60 comparisons often derive from just one or two matched mutant cell lines. We used similar processing steps to derive gene-level mutation data from variant call data for the NCI-60, GDSC, and CCLE (Materials and Methods). Still, differences between the underlying sequencing technologies and initial data preparation methods across the sources may account for some of the observed discrepancies. For example, the CCLE mutation data were obtained for a selected set of 1,667 cancer-associated genes subject to high-depth exome capture sequencing (5), and consistently yielded the largest numbers of cell lines with function-impacting mutations. The greater number of mutations found for *KRAS*, *PTEN*, *BRAF*, *NRAS* or *MSH6* in CCLE relative to the GDSC or NCI-60 databases (evaluated by global exome sequencing) shown in Supplementary Figure 2 reflects the importance of sequencing depth for accurate assessment of mutations.

For a more focused and translational assessment of mutation data concordance, we examined the overlap between sources for established oncogenes and tumor suppressor genes (Figures 2c-d, Supplementary Table 1). For the tumor suppressors, we binarized the data using a probability threshold of 0.7 (to account for the recessive nature of such mutations), while for the oncogenes, the threshold of 0.3 was retained (to account for the dominance of oncogene activating mutations). The most frequently mutated genes were TP53, KRAS, BRAF, APC, RB1, NF1, PTEN, SMARCA4 and MLH1 (Supplementary Table 1). BRAF mutation profiles showed substantial overlap (J > 0.7) across all source comparisons, as was the case for the TP53 gene across the GDSC and CCLE (J = 0.69). On the other hand, PIK3CA, BRCA2, BRCA1 and MLH1, MSH6 and MSH2 mutation comparisons were largely divergent. These discrepancies reflect the ongoing challenges and tradeoffs with mutation profiling technologies and mutation calling procedures. The ability of CellMinerCDB to compare and integrate data across sources highlights the fundamental research efforts and technological standards still required for accurate identification of mutations.

### Drug activity data reproducibility and enrichment

Recent studies have examined the issue of drug data reproducibility, noting potential sources of data divergence such as assay type and duration of drug treatments (13–15). To explore reproducibility and enrich for genomic signatures, we tested a selected set of NCI-60-screened compounds for activity in the larger GDSC panel (Supplementary Table 2). Noting that the GDSC and the NCI/DTP used different assays to determine their IC50 values (Cell Titer Glo measurements of ATP at 72 hours post-treatment versus sulforhodamine B measurement of total protein at 48 hours post-treatment, with additional differences in cell seeding densities and drug dose ranges), we compared 19 drugs referenced by their NSCs (National Service Center identifiers) and associated with a range of mechanisms of action. The drugs with the strongest correlations were bisacodyl and acetalax (Figure 3a, R = 0.84, p = 8.6 × 10^−13^, N = 44, R = 0.80, N = 43, p = 1.0 × 10^−10^, respectively). These FDA-approved laxatives were included in our comparative analysis due to their outstanding activity in two of the three triple-negative breast cancer cell lines of the NCI-60, and the lack of pre-existing data in the CTRP, CCLE and GDSC. The GDSC results confirm that acetalax and bisacodyl elicit a broad range of cytotoxic responses in the expanded GDSC cell line collection, and extend our NCI-60 observations, being more active than any of the 15 tested oncology drugs by a significant margin (p < 7 × 10^−10^) in the 22 GDSC triple-negative breast cancer lines (Supplementary Table 3).

**Figure 3:**
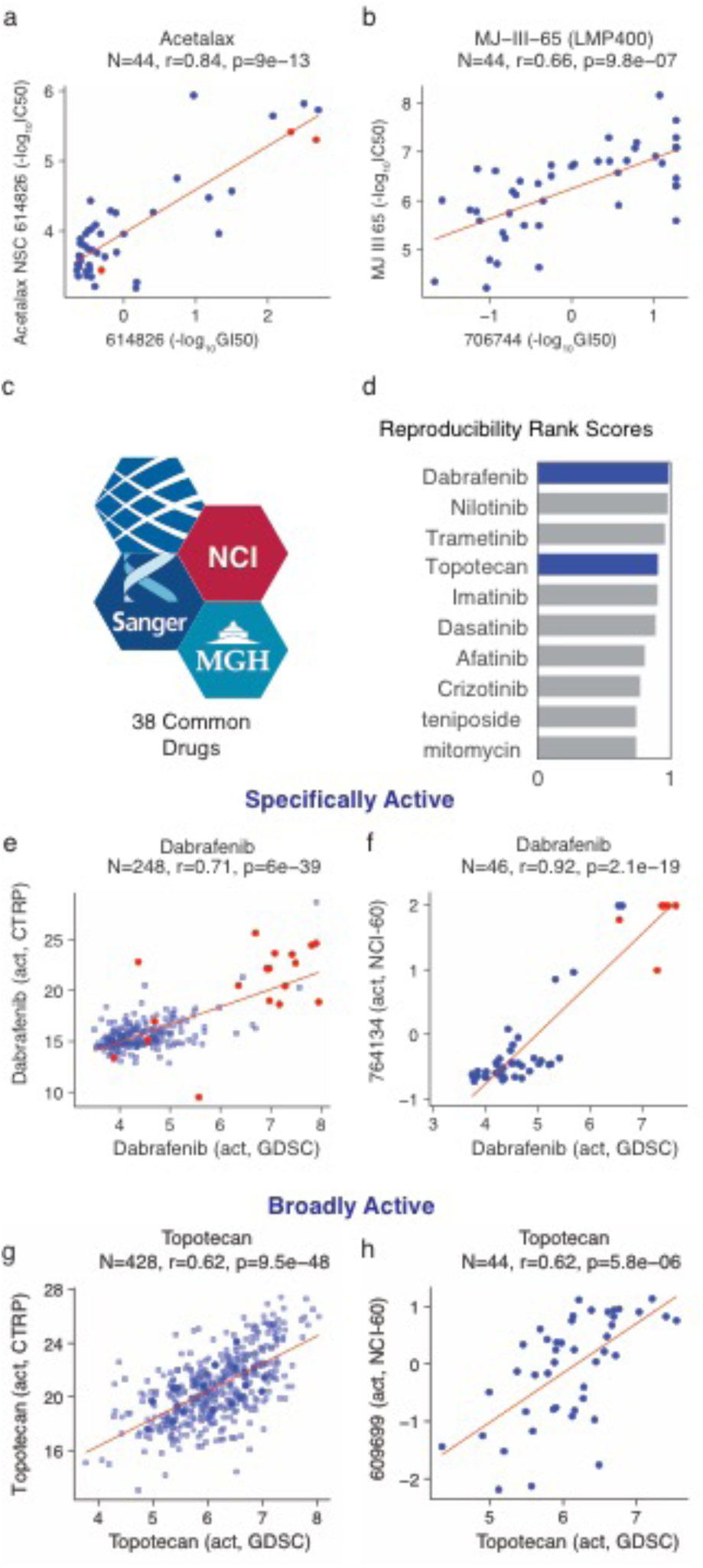
Drug activity data reproducibility. GDSC versus NCI-60 drug activity data in matched cell lines for (a) acetalax and (b) MJ-III-65 (LMP744). Each point represents a matched cell line and red points in (a) indicate triple-negative breast cancer cell lines. (c, d) Ranking of drugs with the best correlated activity data across sources. 38 drugs were tested in the NCI-60, GDSC, and CTRP. For each of the three inter-source comparisons, drugs were ranked by activity correlation strength (q-value), with ranks scaled between 0 (lowest) and 1 (highest). Drugs were then ranked by the average of the three inter-source comparison rank scores. Strong activity correlations occur between specifically active compounds, such as the BRAF inhibitor dabrafenib (e, f), where melanoma lines are shown in red, and broadly active compounds such as the topoisomerase I inhibitor topotecan (g, h). The NCI-60-matched data in (f) and (h) captures the pattern observed with matched data between the larger GDSC and CTRP collections. The full data table excerpted in (d) is shown in Supplementary Figure 6.

The non-camptothecin indenoisoquinoline-based topoisomerase I inhibitor in clinical trial, LMP744 (NSC 706744; MJ-III-65) (16, 17), was also included in our 19-compound test set to assess the similarity of its activity profile with that of topotecan over a larger cell line collection and enrich the genomic signature associated with its activity (R = 0.83, p = 4.2 × 10^−187^, N = 715). LMP744 exhibited solid NCI-60/GDSC activity data concordance (R = 0.66, p = 9.8 × 10^−7^, N = 44) (Figure 3b). Overall, 16 of the 19 tested compounds across the NCI-60 and GDSC gave significant Pearson’s correlations (Supplementary Table 2). Difficulties with the assay were evident for the three drugs with more discrepant activity data. Dacarbazine, an alkylating agent related to temozolomide, and vincristine, an anti-tubulin, both had poor reproducibility even within DTP assay replicates, while the DTP assay for fulvestrant appeared to be out of proper concentration range (Supplementary Figure 5).

**Figure 4:**
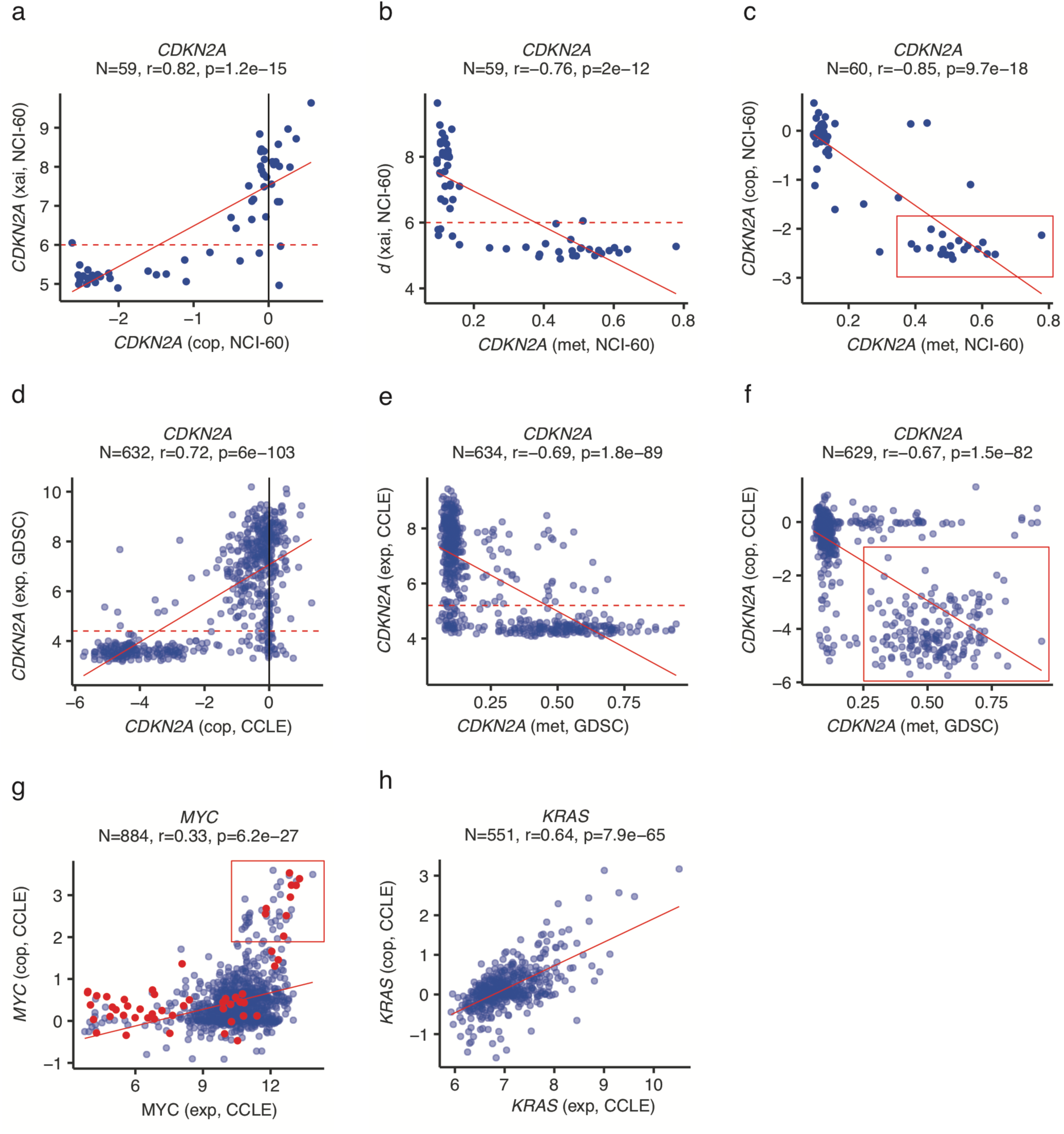
Exploring gene expression determinants. Reduced mRNA expression (xai, average log2 intensity) of the cell cycle inhibitor and tumor suppressor CDKN2A (p16) is associated with DNA copy loss (cop) (a) and promoter methylation (met) (b) in the NCI-60 lines. In a subset of NCI-60 lines, enclosed in the red box, (c), DNA copy loss accompanies higher levels of promoter methylation. DNA copy number and promoter methylation data from the CCLE and GDSC, respectively, can be also be visualized over matched cell lines to verify a similar pattern in larger cell line collections (d-f). DNA copy number gain is associated with increased expression (exp, z-score microarray log2 intensity data) of the oncogenes MYC (g) and KRAS (h) in selected CCLE cell lines. In (g), small cell lung cancer lines are indicated in red to highlight a subset potentially derived from MYC-driven tumors (within red box).

**Figure 5:**
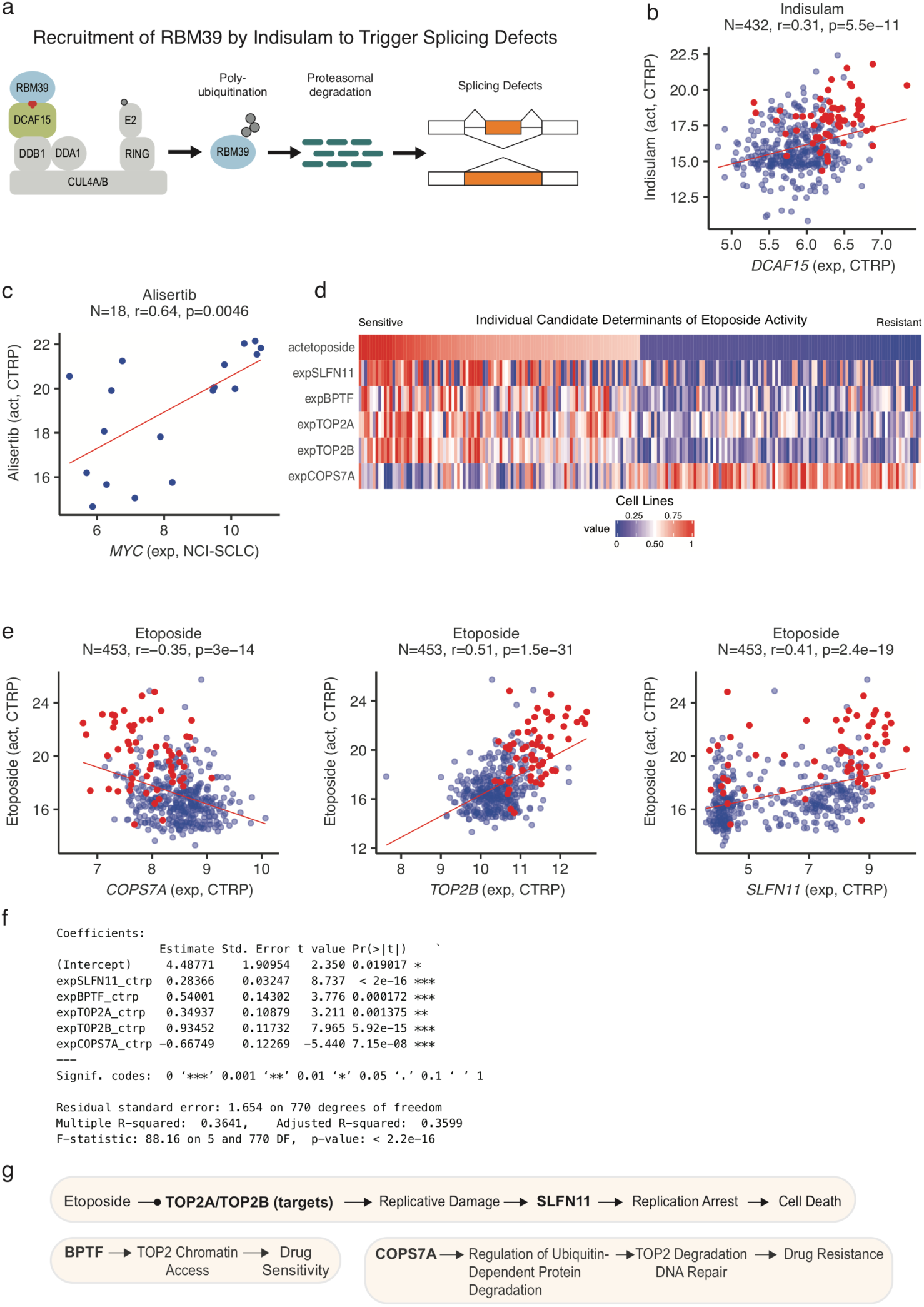
Exploring drug response determinants. (a, b) Response to the pre-mRNA splicing inhibitor indisulam versus expression of its target complex component DCAF15 in the CTRP. Drug response is in (b) is measured by the activity area above the dose-response curve, with higher values indicating relative drug sensitivity. A report of increased indisulam sensitivity in hematopoietic cell lines (shown in red) with high DCAF15 expression is readily verified (20). (c) Response to the Aurora kinase inhibitor alisertib is associated with increased MYC expression in small cell lung cancer lines (21). (d-g) Individual candidate determinants of drug response can be integrated within a multivariate linear model, as shown for etoposide. Additional associations can be identified using partial correlation analyses to select complementary correlates for an existing model. Determinant selection for multivariate linear models can also be performed using the LASSO algorithm.

Further focusing on drug activity data reproducibility, we analyzed the 38 drugs tested in all 3 databases (Figure 3c). CCLE was excluded because of its small drug dataset (24 drugs), which was largely included in CTRP. For each of the three inter-source comparisons, drugs were ranked by activity correlation strength [q-value, scaled between 0 (lowest) and 1 (highest)]. The drugs were then ordered by the average of the 3 inter-source comparison rank scores (Figure 3d, Supplementary Figure 6). As noted in earlier studies of drug activity data reproducibility (15), strong activity correlations were observed for *specifically active* compounds (Figures 3e and 3f), such as the BRAF inhibitor dabrafenib, where outstanding responses occur in cell lines with the activated kinase target. Notably, we also observed high correlations for *broadly active* drugs, such as the topoisomerase I inhibitor topotecan (Figures 3g and 3h), indicating that the cancer cell line responses are reproducible across databases and assays. For many of the 38 assessed drugs (see lower half of Supplementary Figure 6), there were discordant activity data between one or more pairs of sources. The inter-source activity data comparisons of CellMinerCDB allow individual researchers to identify drugs with concordant data, as well as those for which inconsistencies arise, so they can pursue molecular pharmacology analyses only with reliable data.

**Figure 6:**
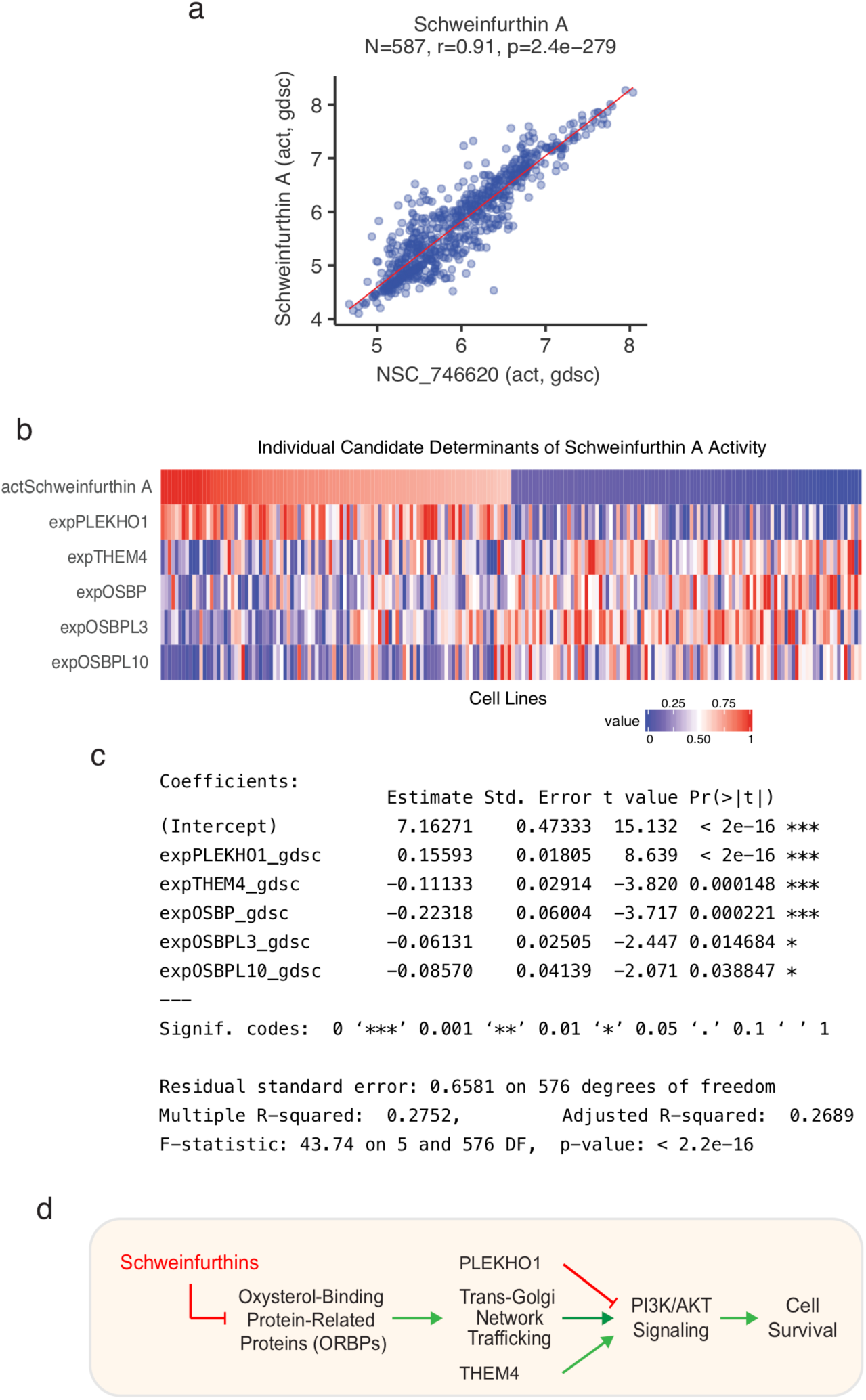
A multivariate model of Schweinfurthin A drug activity. (a) Reproducibility of the data for the two schweinfurthin derivatives tested in the GDSC. (b, c) Integration of molecular correlates of Schweinfurthin activity within a multivariate linear model. The heatmap in (a) shows drug activity and gene expression data for the 100 most sensitive (left) and resistant (right) GDSC cell lines, with higher values in red and lower values in blue. (d) Scheme of the proposed molecular pharmacology of the schweinfurthins. Schweinfurthins have been shown to inhibit PI3K/AKT signaling and cell survival by binding oxysterol binding protein-related proteins (ORPs) to disrupt trans-Golgi-network trafficking required for robust pathway activity (28). Together with the ORPs OSBP, OSBPL3, and OSBPL10, the other candidate determinants, PLEKHO1 and THEM4, have also been implicated in PI3K/AKT signaling (27, 31). Plots and analyses in panels b - d are based on non-hematopoietic GDSC cell lines.

### Exploring gene regulatory determinants

Cancer-specific gene expression is often altered by promoter methylation or DNA copy number changes. CellMinerCDB makes it easy to identify these and other potential gene regulatory determinants. For example, in the NCI-60, reduced expression of the tumor suppressor gene *CDKN2A* (p16^INK4a^) is associated with both DNA copy loss (Figure 4a) and promoter methylation (Figure 4b) across diverse tissue types. Furthermore, Figure 4c shows that approximately 25% of NCI-60 cell lines show both alterations, consistent with bi-allelic, ‘two-hit’ suppression of *CDKN2A* expression. Integration of matched cell line GDSC methylation data and CCLE DNA copy number data illustrates the same *CDKN2A* regulatory relationships in a larger cell line collection (Figures 4d-f). *CDKN2A* is unique with respect to the high proportion of cell lines showing co-occurrence of promoter methylation and DNA copy loss (Supplementary Table 4). The impact of copy gain on increased oncogene expression can be similarly assessed with CellMinerCDB. Figure 4g shows that a subset of *MYC*-driven CCLE small cell lung cancer lines exhibit both *MYC* copy gain and increased *MYC* gene expression. KRAS activation, typically regarded as mutation-driven, also occur by copy gain, as evidenced in a subset of CCLE lines (Figure 4h), as well as in clinical studies (18).

### Exploring drug response determinants

CellMinerCDB provides correlation analyses and scatter plots for testing and visualizing potential response-determinant relationships (univariate analyses) as well as multivariate linear regression methods for integrating multiple determinants (multivariate analyses; see Figures 5d and 6b). CellMinerCDB also retrieves significant candidate genomic determinants as well as drug correlations (compare analysis tool). This type of approach led to the discovery of Schlafen 11 (*SLFN11*), a new molecular determinant of response to a broad range of widely used DNA-targeted anticancer agents including topoisomerase inhibitors, platinum derivatives, PARP inhibitors and antimetabolites (5, 19). CellMinerCDB allows users to quickly assess the generality of results presented in the literature, and iteratively explore evidence for multifactorial mechanistic models. Figure 5a shows an example for indisulam, which targets the splicing factor RBM39 for proteasomal degradation by forming a ternary complex with RBM39 and the E3 ubiquitin ligase receptor DCAF15. A report of increased indisulam sensitivity in hematopoietic cell lines with high *DCAF15* expression is readily verified with CellMinerCDB (Figure 5b, red dots) (20). CellMinerCDB-integrated pharmacogenomic data also corroborate a report of *MYC*-driven small cell lung cancer exhibiting vulnerability to aurora kinase inhibition (21) (Figure 5c).

Multivariate model development without bioinformatics assistance is illustrated for the topoisomerase II inhibitor etoposide (Figure 5d-g). Starting with expression of the drug target (*TOP2A* and *TOP2B*) (22) and *SLFN11*, an established determinant of response to DNA-targeted drugs (19,23,24), the user can select additional determinants based on biological knowledge. Determinant selection can be further guided by pathway annotations and partial correlation analyses, which rank additional features by their capacity to improve the current model (Figure 5g).

### A multivariate model of Schweinfurthin A drug activity

Schweinfurthin A was discovered through an NCI natural products initiative aimed at identifying novel anticancer compounds with distinctive NCI-60 activity profiles indicative of a novel target [COMPARE analysis (25)]. Its wide activity range, with notable potency in leukemia and CNS lines (< 10 nmol/L), motivated synthesis of a series of derivatives, which retained the same unique activity profile in the NCI-60 (26). Further development of schweinfuthins has been hampered by lack of understanding of their molecular target. We tested Schweinfurthin A and 5’-methylschweinfurthin G on approximately 600 cell lines of the GDSC panel and applied the various features of CellMinerCDB to reveal the molecular pathways for drug response. The activity of the two schweinfurthin compounds was strongly correlated over the GDSC lines (R = 0.87, p = 8.8 × 10^−182^, N = 585, Figure 6a). The CellMinerCDB Univariate Analysis tool (Compare Patterns tab) indicated that one of the strongest molecular correlates of schweinfurthin activity is expression of *PLEKHO1*, a negative regulator of PI3K/AKT signaling (R = 0.47, p = 1.95 × 10^−33^, N = 582) (27). This result is consistent with a recent study showing that schweinfurthins inhibit mTOR/AKT signaling by interfering with trans-Golgi-network trafficking (TGN) (28). In particular, schweinfurthins bind to oxysterol-binding proteins that regulate TGN trafficking (29, 30), thereby arresting lipid raft-mediated PI3K activation and functional mTOR/RheB complex formation.

Using the multivariate analysis feature of CellMinerCDB (“Regression Models” tab; https://discover.nci.nih.gov/cellminercdb/), we developed a linear predictive model for schweinfurthin response integrating expression of *PLEKHO1*, *THEM4*, a positive regulator of AKT signaling (31), and the genes encoding the oxysterol binding protein family members *OSBP*, *OSBPL3*, and *OSBP10* (Figure 6c-d). Increased expression of the latter conceivably sustains TGN trafficking and PI3K/AKT signaling, in keeping with their negative regression coefficient weights as resistance determinants in the model. The negative weighting of *THEM4* expression and positive weighting of *PLEKHO1* expression are similarly consistent with their respective roles in activating and suppressing PI3K/AKT signaling. These analyses give molecular insight into the cholesterol trafficking and intracellular membrane pathways as the targets of sweinfurthins, and open new opportunities for matching the activity of schweinfurthins with genomic and molecular signatures.

### Relating EMT status with gene expression to identify a novel EMT gene and schweinfurthin activity in mesenchymal cells

Epithelial-mesenchymal transition (EMT) is a fundamental process in development, wound healing and cancer progression, characterized by the loss of cell-cell adhesion and the acquisition of motile and invasive properties (Figure 7a) (32, 33). EMT is driven by dominant transcription factors, including ZEB1/2, SNAI1/2, and TWIST1/2, and is reversible through a continuum of states from epithelial to mesenchymal. These attributes have motivated the development of gene expression-based EMT signatures to identify cell line state and understand drug resistance.

**Figure 7:**
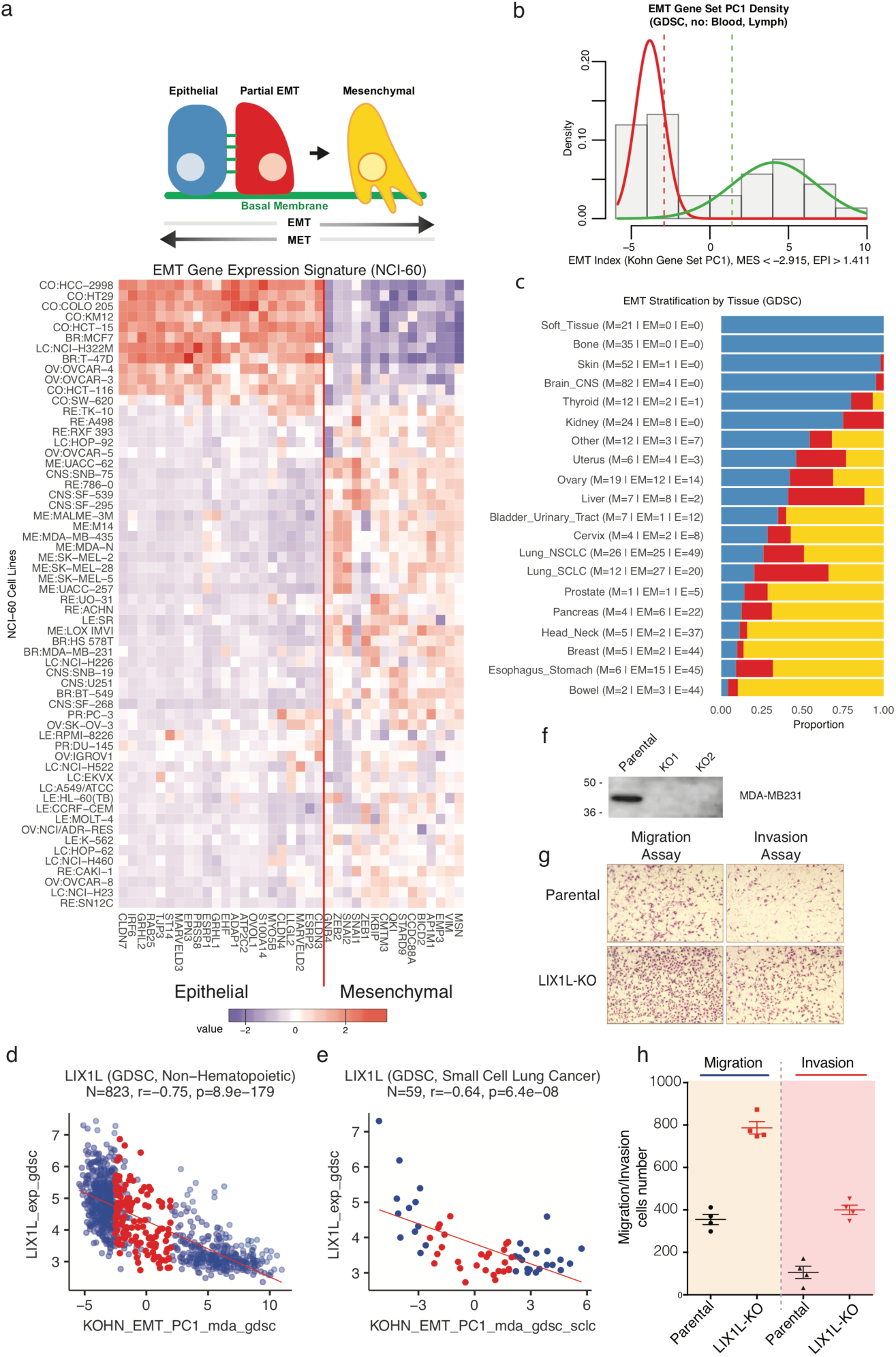
Relating Epithelial Mesenchymal Transition (EMT) status with gene expression to identify *LIX1L* as a novel EMT gene. (a) A 37 gene EMT signature developed in (34) was used to derive a numerical index of EMT status as a weighted sum of cell line-specific EMT gene expression values (see Supplementary Methods for details). Epithelial and mesenchymal status is associated with positive and negative index values, respectively. (b) For 821 non-hematopoietic cell lines in the GDSC collection, the EMT index values show a bimodal distribution, which can be modeled as a Normal mixture. Cell lines with EMT index values less than (greater than) one standard deviation above (below) the putative mesenchymal (epithelial) group mean are annotated as mesenchymal (epithelial). (c) EMT Stratification by tissue of origin. (d, e) Expression of *LIX1L*, a novel mesenchymal gene, is strongly correlated with the EMT index signature. ‘Epithelial-Mesenchymal’ lines with intermediate EMT index values are indicated in red. Mesenchymal lines are in blue at left, and epithelial in blue at right. (f - h) Knockout of *LIX1L* in the MDA-MB231 cell line results in increased cell migration and invasion.

We applied a 37 gene EMT signature initially developed in the NCI-60 (34) to derive a numerical index of EMT status as a weighted sum of cell line-specific EMT expression values (Supplementary Methods). Figure 7b shows a bimodal distribution for the EMT index values of the 821 non-hematopoietic GDSC cell lines, allowing cell lines to be stratified into epithelial, mesenchymal, and epithelial-mesenchymal categories. EMT stratification within particular tissues of origin showed a substantial proportion of intermediate epithelial-mesenchymal lines in several tissue types, including liver, ovary, and lung (Figure 7c). The numerical EMT index is available for all CellMinerCDB-integrated data sources as the variable KOHN_EMT_PC1 (‘Metadata’), allowing its correlations with molecular and drug response features.

Using our numerical EMT index, we identified *LIX1L*, as a novel mesenchymal gene whose expression is highly correlated with the EMT index signature (34) across multiple cancer types (Figure 7d, R = -0.75, p = 8.9 × 10^−179^, N = 823). *LIX1L* is also broadly expressed in TCGA tumor samples (Supplementary Figures 3a). Its expression is not correlated with its homolog *LIX1* (Supplementary Figures 3b-d). Experimental analyses in knockout cell lines revealed that *LIX1L* is inversely connected with cell migration and invasiveness (Figure 7f-h, Supplementary Figures 3c, d).

We also correlated the EMT index with GDSC activity profiles for 297 compounds (Supplementary Figure 4). Schweinfurthin A emerged as the strongest correlate, indicating relatively increased activity in mesenchymal cancer cell lines, such as those from derived from bone or soft tissue (Supplementary Figure 4b). The second strongest negative correlate with the EMT index was the RHO-associated kinase 1 inhibitor GSK269962A (Supplementary Figure 4b), whose target participates in the regulation of actin dynamics and cell motility during the EMT (32, 33).

## Discussion

CellMinerCDB allows researchers to interact freely with an unparalleled breadth of cancer cell line pharmacogenomic data, using their expertise to iteratively refine analyses. Our results, spanning data assessment, integration, and discovery demonstrate the value of working across data sources. They support the maturity and essential reproducibility of most molecular profiling technology platforms, such as gene transcript expression, copy number, and methylation.

Mutation data are prominently featured in translational studies, and CellMinerCDB exposes the issue of discrepancies between matched cell line mutation profiles across sources. This provides a foundation for understanding and mitigating sources of variability, which reflect the ongoing technical challenges and tradeoffs with acquiring genome sequencing data. Somatic variant calling in cancer cell lines is inherently challenging due to the absence of matched normal tissue for comparison, as well as the potentially higher mutation burden in cell lines relative to primary tumor tissue. In this setting, variability in cell line mutation data across sources may additionally arise from differences in variant calling algorithms, as well as data sources used for filtering likely germline variants. One approach for excluding potentially cell line-specific passenger mutations is to filter variants based on frequency in patient populations (4). Another strategy for more robust mutation data is to pursue higher depth targeted sequencing of a restricted gene set. Indeed, we noted that CCLE data, derived from the latter sort of approach, consistently identified more mutant cell lines for prominent oncogenes and tumor suppressor genes than genome-wide exome sequencing (as in the NCI-60 and GDSC databases). Nevertheless, using CellMinerCDB, researchers can quickly integrate mutation data across sources to identify cell lines with consistent mutation calls for a gene of interest, together with other potentially impacted lines.

Drug response assays used across the major pharmacogenomic data sources measure different biochemical features over different time scales. Still, CellMinerCDB demonstrates significant concordance between activity data generated at the NCI and the Sanger Institute. Moreover, for several widely used anticancer drugs, such as topotecan and dabrafenib, we observed activity data reproducibility across all major sources. We expanded the data available for understanding variability in drug activity profiling by testing 19 compounds from a range of mechanistic classes in the large GDSC cell line panel. Readily accessible scatterplots of matched cell line activity data can highlight problem areas with particular assays, such as inappropriate concentration ranges in the case of fulvestrant in the NCI/DTP assay. The cross-database comparison features of CellMinerCDB allow researchers to explore these issues and focus on drugs with reliable data. Among the 19 drugs tested, we found consistent results for most of the drugs and identified bisacodyl and acetalax as potential drug repurposing candidates, being substantially more active in triple-negative breast cancer lines relative to other cancer drugs tested on the GDSC panel.

With both drug activity reproducibility and broader associations between molecular features, such as *CDKN2A* expression and gene copy/methylation, we noted that the NCI-60 could effectively capture relationships evident in larger cell line sets. The latter better reflects tissue type diversity and associated context-specific molecular features. Still, for dominant associations, such as *SLFN11* expression and DNA-targeted drug responses, representative cell line sets such as the NCI-60 are often sufficient (19). The NCI-60 provides drug responses for over 21,000 individual agents, making it an unmatched resource for discovery of new chemotypes based on correlations with genomic data and response patterns to drugs with known targets. The NCI databases are also a tractable starting point for molecular data expansion with leading-edge technologies. RNA-Seq data with isoform-specific transcript expression, and SWATH mass spectrometry-based protein expression data have been generated for the NCI-60, and made available within CellMinerCDB (35).

CellMinerCDB ultimately aims to provide a seamless platform, integrating not just data sources, but the intuition and expertise of experimental scientists and clinicians. The present publication provides only a sample of the potential of CellMinerCDB for focusing translational studies. CellMinerCDB uniquely complements existing data portals that provide detailed information on their associated data, together with specialized analyses. Each source offers unique strengths; for the NCI-60 a broad array of molecular profiling data and the largest set of tested compounds, including potentially novel molecules for drug development, such as the indenoisoquinoline topoisomerase I inhibitors (17); for the GDSC, a growing range of molecular and drug activity data over a large cell line panel; for the CTRP, the largest set of tested compounds in a large cell line panel; for the CCLE, high-depth mutation profiling and other molecular data, readily extensible to the CTRP via CellMinerCDB. By empowering researchers to easily build on these strengths and pursue their own questions, CellMinerCDB will advance the potential of cancer cell line pharmacogenomic data to lay the foundation, validate, and focus experimental and ultimately clinical studies.

## Methods

### CellMinerCDB Implementation

CellMinerCDB was implemented using the R programming language, with interactive features developed using RStudio’s Shiny web application framework (https://shiny.rstudio.com/). Underlying analyses and data representations were built with functionality provided by our publicly available rcellminer R/Bioconductor package (8). For each data source, R data packages were constructed, using software components defined within rcellminer to integrate drug activity data, molecular profiling data, and associated cell line, drug, and gene annotations. This standard data representation allowed diverse data sources to be readily integrated within CellMinerCDB. Source-specific data used in CellMinerCDB data package construction are described in the sections below.

### NCI-60 Data

NCI-60 drug activity, molecular profiling, and annotation data was obtained from CellMiner (Database Version 2.1). The latest versions of these data can also be downloaded from https://discover.nci.nih.gov/cellminer/loadDownload.do. Detailed information is provided in (1,9,10,36). Essential attributes made available within CellMinerCDB are summarized below.

#### Compound activity

Standardized, ‘z-score’ values were derived from measurement of 50% growth-inhibitory (GI50) concentrations using the sulforhodamine B total protein cytotoxicity assay. For each compound, the mean and standard deviation of -log10[molar GI50] values over the NCI-60 lines are used to center and scale the data.

#### Gene expression

Integration of relevant probe-level data from 5 microarray platforms (1) is provided in both standardized ‘z-score’ form, derived as described above for the drug activity data, and as average log2 intensities.

#### Gene-level mutation

The mutation data value for a given gene and cell line is derived from computed probability of a homozygous function-impacting mutation, which is then expressed as a percentage. NCI-60 exome sequencing data was obtained and processed as described (9). Missense mutations were functionally categorized using ANNOVAR (37). Missense mutations with a frequency > 0.005 in either the ESP6500 or 1000 Genomes normal population datasets (i.e., potential germline variants) were excluded, together with mutations predicted not to impact protein function by the SIFT and PolyPhen2 algorithms (SIFT > 0.05 or PolyPhen2 HDIV < 0.85 or PolyPhen2 HVAR < 0.85). To obtain a summarized, gene-specific mutation value for each cell line, the probability of both alleles having at least one of the variants was computed. Specifically, let x = (x_1_, …, x_n_) be a vector of gene-associated mutation conversion fraction values for a given cell line. The summary gene mutation probability value for this cell line is computed as 1 – (1 – x_1_) … (1 – x_n_), and then converted to a percentage value.

#### DNA copy number

DNA copy data were integrated from four array-CGH platforms (36). Numerical values indicate the average log2 probe intensity ratio for the cell line (gene-specific chromosomal segment) DNA relative to normal DNA.

#### DNA methylation

Data were obtained using the Illumina Infinium Human Methylation 450 platform as described (10).Values lie between 0 (lack of methylation) and 1 (complete methylation).

#### microRNA expression

Data were obtained using the Agilent Technologies Human miRNA Microarray V2 (38). Numerical values indicate average log2 probe intensity.

#### Protein expression

Reverse phase protein array (RPPA) data were obtained as described (39). Numerical values indicate probe intensities.

### GDSC Data

#### Compound activity

Preprocessed activity data for 256 compounds were downloaded from http://www.cancerrxgene.org/downloads. GDSC-provided activity values were converted to indicate the - log10[molar IC50].

#### Gene expression

Raw Affymetrix Human Genome U219 microarray data deposited in ArrayExpress (E-MTAB-3610) were processed using RMA normalization. Probe-to-gene mapping was performed using the BrainArray CDF file for the Affymetrix HG-U219 platform, available at http://brainarray.mbni.med.umich.edu/Brainarray/Database/CustomCDF/17.1.0/entrezg.download/HGU2_19_Hs_ENTREZG_17.1.0.zip. Numerical values summarize gene-specific log2 probe intensities. Additional platform and processing details are provided in (4).

#### Gene-level mutation

A tab-separated table listing variants detected in GDSC cell lines was downloaded from COSMIC (release v79). Variants indicated as heterozygous and homozygous were assigned values of 0.5 and 1, respectively. After this, gene-level mutation values were computed as described for the NCI-60 mutation data, except that the final, gene and cell line-specific mutation probabilities were retained (rather than converted to percentage values).

#### DNA methylation

The table of pre-processed beta values for all CpG islands across the GDSC cell lines was downloaded from the supplementary resources site http://www.cancerrxgene.org/gdsc1000/GDSC1000_WebResources/ (4). DNA methylation data were obtained using the Illumina Infinium Human Methylation 450 platform, and gene-level methylation values were computed using the approach utilized with the NCI-60 data (10).

#### Determination of prospective triple negative breast cancers

Expression levels of *ERBB2*, *ESR1*, *ESR2* and *PGR* were assessed by GDSC using the Affymetrix Human Genome U219 Array and accessed in CellMinerCDB. Cell lines with a low value for all 3 genes were classified as triple negative. The log2 intensity thresholds used were *ERBB2*<5, *ESR1*< 3.5, and *PGR*<3.

### CCLE Data

CCLE data were downloaded from https://portals.broadinstitute.org/ccle/home (5).

#### Compound activity

Activity profiles are available for 24 compounds. CCLE-provided activity values were converted to indicate the -log10[molar IC50].

#### Gene expression

Raw CEL file data derived from the Affymetrix U133+2 platform were downloaded from the CCLE portal. Normalization was performed using the frma method, implemented by the corresponding Bioconductor package (40). Numerical values are the average of gene-specific log2 probe intensities, with the gene-to-probe-set mapping obtained from the hgu133plus2.db Bioconductor package.

#### Gene-level mutation

The table of targeted sequencing-based mutation data for 1651 genes was downloaded from the CCLE portal. Using the provided allelic fraction information for individual variants, gene-level mutation values were computed as described for the NCI-60.

#### DNA copy number

Data derived from the Affymetrix SNP 6.0 array were downloaded from the CCLE portal. Numerical values are normalized log2 ratios, i.e., log2(CN/2), where CN is the estimated copy number.

### CTRP Data

Activity data for 481 compounds across 823 cell lines were obtained from Supplementary Tables S2, S3, and S4 of reference (6). Activity data originally indicated as the area under a 16-point dose response curve (AUC) were subtracted from the maximum observed AUC value (over all cell lines and drugs) to represent activity by the estimated area above the dose-response curve. This transformation allows increased drug sensitivity to be associated with larger values of the activity measure, consistent with other source activity data integrated within CellMinerCDB. The above CTRP cell line set is included in the CCLE, and CCLE molecular data are thus used for CTRP analyses in CellMinerCDB.

### NCI-SCLC Data

Compound activity and transcript expression data for the NCI-SCLC data set were downloaded from https://sclccelllines-dev.cancer.gov/sclc/downloads.xhtml. Activity values were converted to indicate the - log10[molar IC50]. Transcript expression values are derived from log2 microarray probe intensities.

### CellMinerCDB Analyses

Data types across sets of cell lines can be plotted with respect to one another within the ‘Univariate Analyses - Plot Data’ tab. From the ‘Univariate Analyses - Compare Patterns’ tab, additional molecular and drug response correlates can be tabulated, with respect to either the plotted x-axis or y-axis variable. Pearson’s correlations are provided, with reported p-values not adjusted for multiple comparisons. The ‘Regression Models’ tab set allows construction and assessment of multivariate linear models. The response variable can be set to any data source-provided feature (e.g., a drug response or gene expression profile across cell lines). Basic linear regression models are implemented using the R stats package lm() function, while lasso (penalized linear regression models) are implemented using the glmnet R package (41). The lasso performs both variable selection and linear model coefficient fitting (42). The lasso lambda parameter controls the tradeoff between model fit and variable set size. Lambda is set to the value giving the minimum error with 10-fold cross-validation. For either standard linear regression or LASSO models, 10-fold cross validation is applied to fit model coefficients and predict response, while withholding portions of the data to better estimate robustness. The plot of cross-validation-predicted vs. actual response values can also be viewed within CellMinerCDB, to assess model generalization beyond the training data.

Additional predictive variables for a multivariate linear model can be selected using the results provided within the ‘Regression Models - Partial Correlation’ tab. Conceptually, the aim is to identify variables that are independently correlated with the response variable, after accounting for the influence of the existing predictor set. Computationally, a linear model is fit, with respect to the existing predictor set, for both the response variable and each candidate predictor variable. The partial correlation is then computed as the Pearson’s correlation between the resulting pairs of model residual vectors (which capture the variation not explained by the existing predictor set). The p-values reported for the correlation and linear modeling analyses assume multivariate normal data. The two-variable plot feature of CellMinerCDB allows informal assessment of this assumption, with clear indication of outlying observations. The reported p-values are less reliable as the data deviate from multivariate normality.

### Metadata

Cell lines of particular tissue or tumor types can be highlighted in two-variable plots. In addition, correlation and regression analyses can be restricted to cell line subsets by either inclusion or exclusion of selected tissue or tumor types. To enable this, all cell lines across data sources were mapped to the four-level OncoTree cancer tissue type hierarchy developed at Memorial Sloan-Kettering Cancer Center (http://www.cbioportal.org/oncotree/). Every cell line has an OncoTree top level specification, such as ‘Lung’, indicating its tissue of origin. Additional OncoTree levels provide more detailed annotation, distinguishing, for example, small cell lung cancer and various types of non-small cell lung cancer. Within the ‘Regression Models’ tab set, LASSO and partial correlation analyses can be restricted to gene sets curated by the NCI/DTB Genomics and Bioinformatics Group.

## Acknowledgements

We would like to thank Dr. David Goldstein of the NCI Office of Science and Technology Resources for supporting the purchase of software required to enable efficient, multi-user access to the CellMinerCDB site.

## Author Contributions

(conception, design, and development) V.N.R., A.L., F.E., L.L., W.C.R., Y.P. (acquisition and preparation of data) S.V., M.S., F.I., C.H.B, M.J.G., W.C.R. (analysis and interpretation of data; development of results) V.N.R., A.L., M.Y., S.V., F.G.S., M.A., A.T., K.K. C.H.B., M.G., W.C.R., Y.P. (experimental validation studies) M.Y. (writing, review, and revision of the manuscript) V.N.R., A.L., M.Y., F.I., F.G.S., M.A., A.T., C.S., C.H.B., M.G., W.C.R., Y.P. (study supervision) W.C.R., Y.P.

## Supplementary Methods

**Supplementary Figure S1:** Inter-source data reproducibility examples for selected genes and molecular data types.

**Supplementary Figure S2:** CCLE vs. GDSC mutant cell line counts for selected oncogenes and tumor suppressor genes.

**Supplementary Figure S3:** Additional LIX1L experimental data.

**Supplementary Figure S4:** Schweinfurthin activity and EMT index correlations.

**Supplementary Figure S5:** GDSC versus NCI-60 drug activity for fulvestrant, indicating inappropriate drug concentration range in NCI-60 activity assay.

**Supplementary Figure S6:** Overall drug activity data reproducibility rankings for 38 compounds tested in the NCI-60, GDSC, and CTRP, integrating pairwise activity correlations between the sources.

**Supplementary Table 1:** Oncogene and tumor suppressor gene mutation call frequencies and overlaps across data sources.

**Supplementary Table 2:** Comparison of drug activities as measured by the GDSC and the NCI/DTP.

**Supplementary Table 3:** Activity of bisacodyl and acetalax in GDSC triple-negative breast cancer lines.

**Supplementary Table 4:** Co-occurrence of gene promoter methylation and DNA copy loss.

### Filtering of gene-level molecular profiling data for inter-source reproducibility analyses

In pairwise (source A vs. source B) comparisons of gene expression and methylation data, genes which were essentially not expressed or methylated in the inter-source matched cell line set were excluded from correlation analyses (since these cases, the latter would be over noisy data near technical detection thresholds). In particular, in inter-source transcript expression data comparisons, we excluded genes for which the 90th percentile expression value, across matched cell lines from both compared sources, was below 6 (microarray, log2 intensity). Similarly, in the methylation data comparisons, genes for which the corresponding 90th percentile methylation value was below 0.3 (average probe beta value) were excluded.

### Derivation of gene expression-based Epithelial-Mesenchymal Transition (EMT) index and cell line stratification

For each data source, the following steps were taken to obtain a numerical measure of EMT status.

1. Microarray expression data (log2 intensity) over non-hematopoietic cell lines were selected for a subset of EMT genes identified in (30); these included 22 epithelial genes (ADAP1, ATP2C2, CLDN3, CLDN4, CLDN7, EHF, EPN3, ESRP1, ESRP2, GRHL1, GRHL2, IRF6, LLGL2, MARVELD2, MARVELD3, MYO5B, OVOL1, PRSS8, RAB25, S100A14, ST14, TJP3) and 15 mesenchymal genes (AP1M1, BICD2, CCDC88A, CMTM3, EMP3, GNB4, IKBIP, MSN, QKI, SNAI1, SNAI2, STARD9, VIM, ZEB1, ZEB2).
2. Data for each gene was centered and scaled by subtracting the mean expression value over the cell line set and then dividing by the corresponding standard deviation.
3. A principal component analysis was performed, with the EMT index obtained as the first principal component.

For a given cell line, the described EMT index is a weighted sum of EMT gene expression values. For all data sources, mesenchymal gene expression values are associated with negative weights, while epithelial gene expression values are associated with positive weights. EMT index values for non-hematopoietic cell lines in each data source show a bimodal distribution (as in Figure 8b), with putative mesenchymal and epithelial lines having negative and positive index values, respectively. The mixtools R package function nomalmixEM was used to fit a 2-component Gaussian mixture model using the source-specific EMT index data. Cell lines with EMT index values less than (greater than) one standard deviation above (below) the putative mesenchymal (epithelial) group mean are annotated as mesenchymal (epithelial); the remaining non-hematopoietic lines are classified as epithelial-mesenchymal. Hematopoietic cell lines were excluded from EMT index value computations and associated classifications.

**Supplementary Figure S1:**
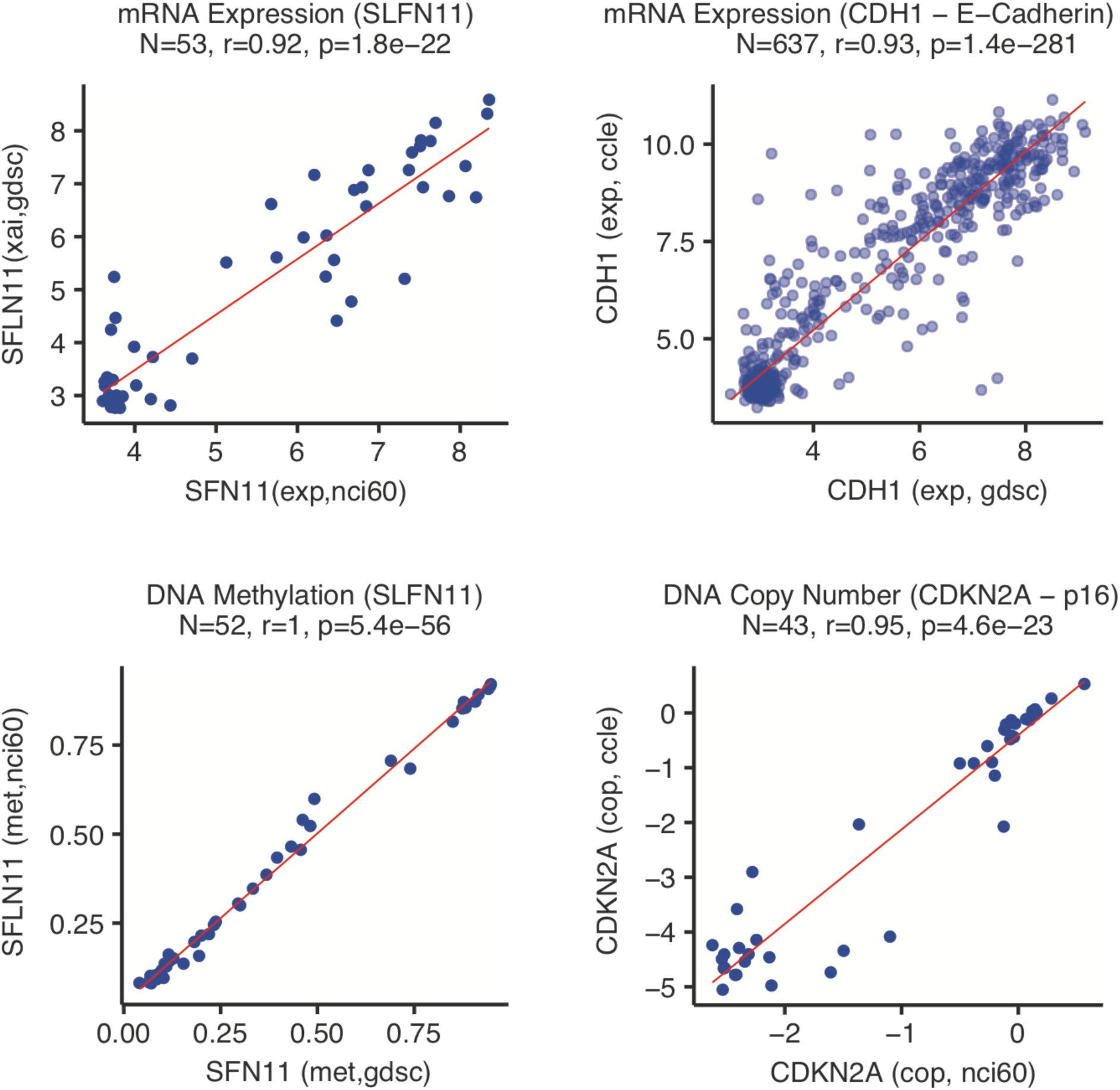
Inter-source data reproducibility examples for selected genes and molecular data types.

**Supplementary Figure S2:**
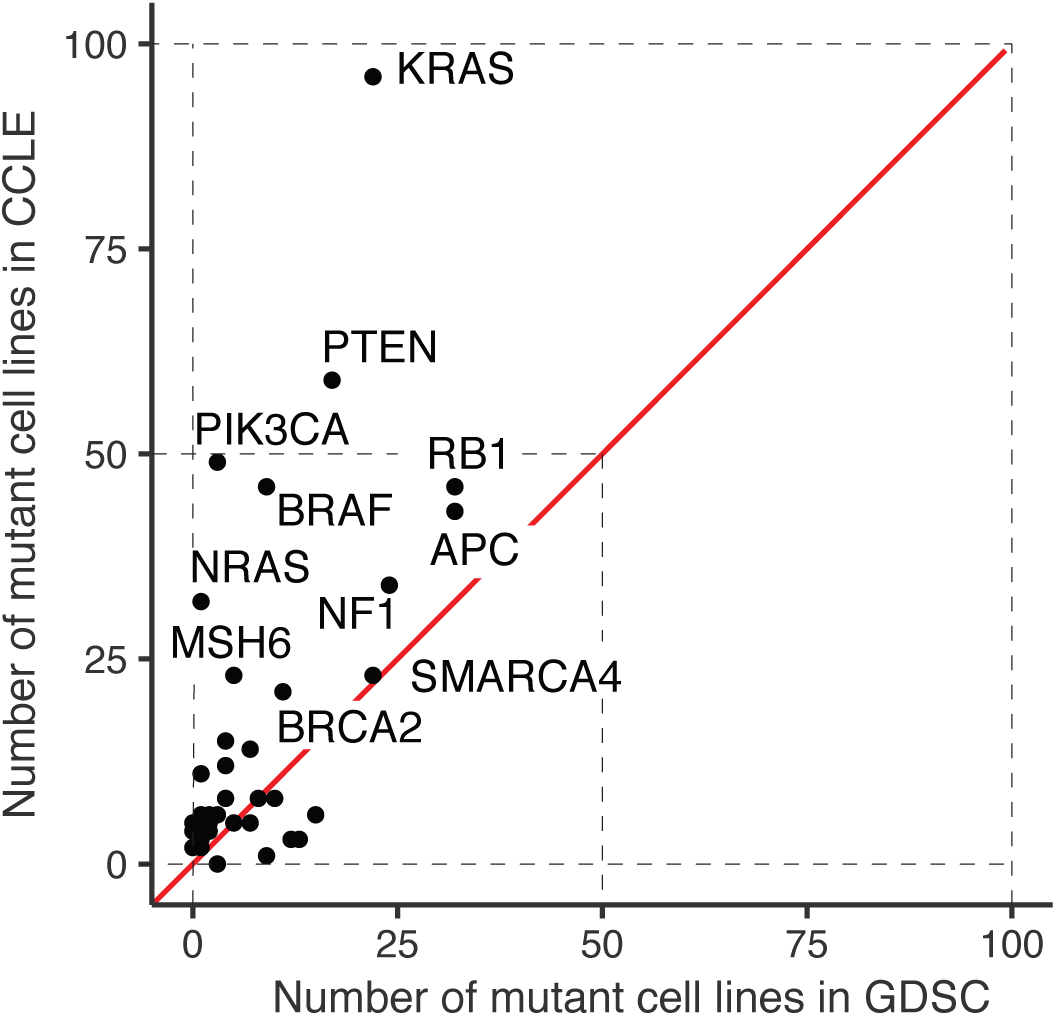
Importance of sequencing depth for retrieving mutant cell lines. CCLE vs. GDSC mutant cell line counts for selected oncogenes and tumor suppressor genes.

**Supplementary Figure S3:**
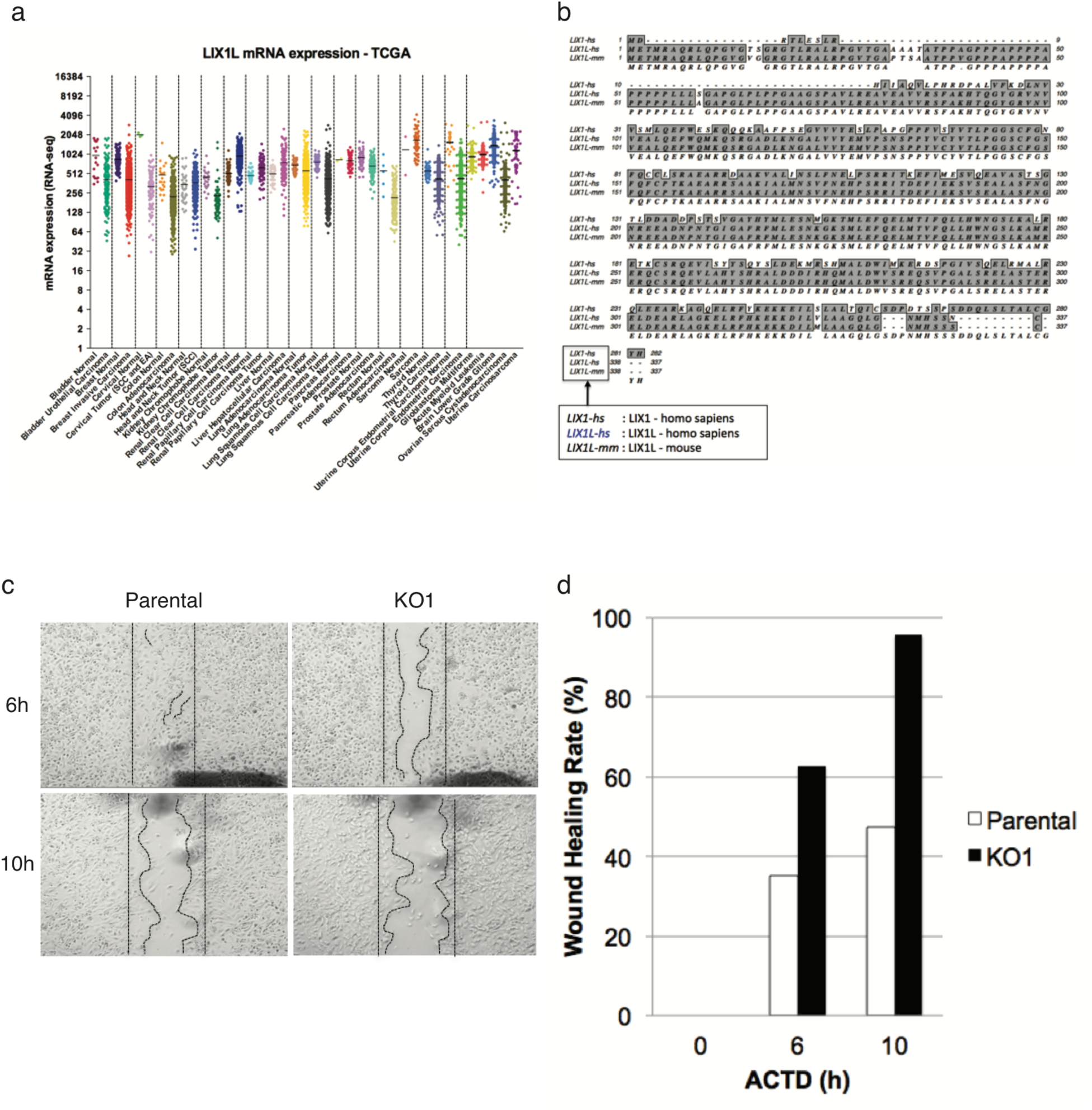
(a) LIX1L transcript expression across TCGA tumor samples. (b) Sequence alignment showing homology between LIX1, LIX1L (human), LIX1L (mouse). (c, d) Scratch-wound assay results showing increased cell migration with LIX1L knockout.

**Supplementary Figure S4:**
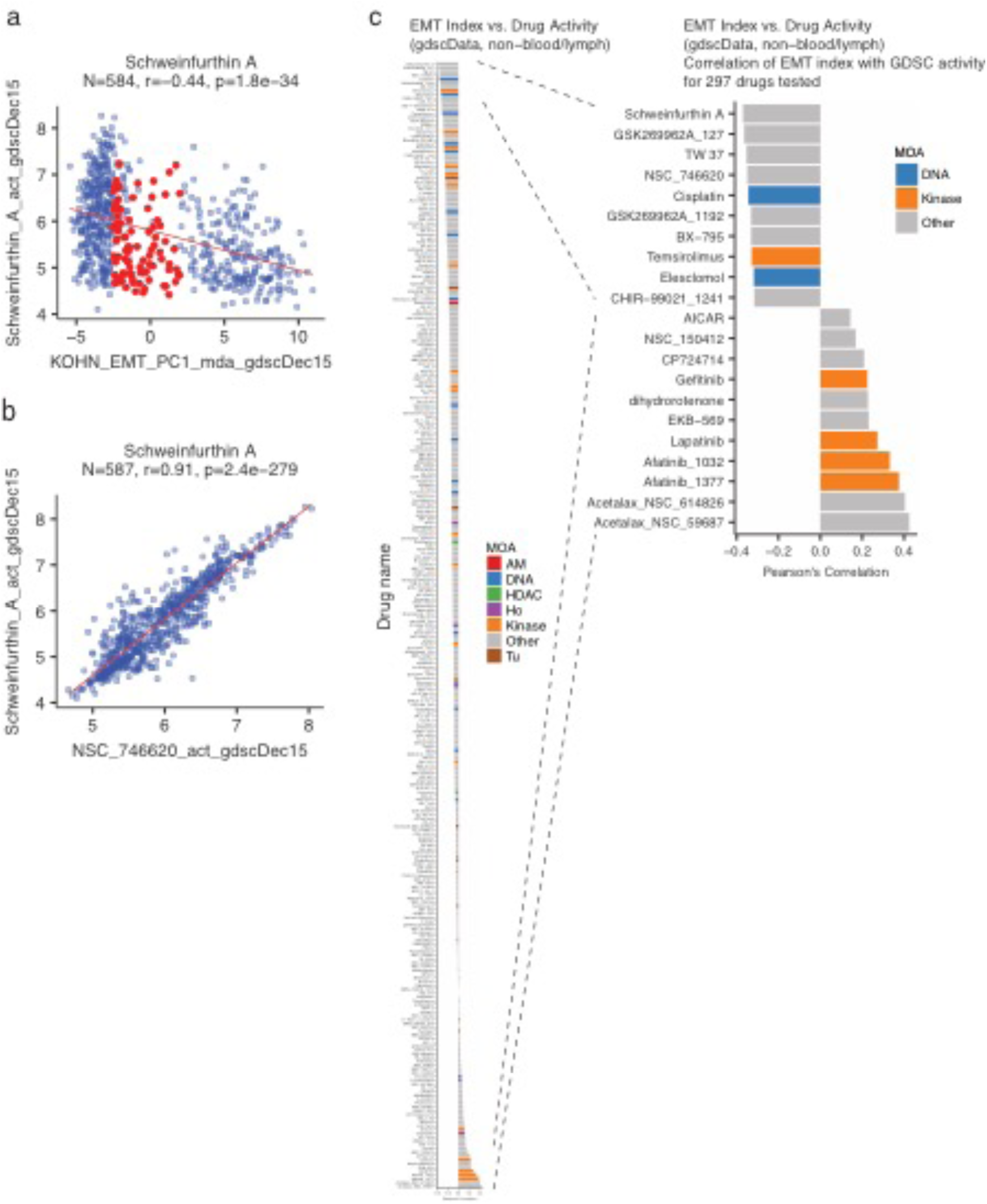
(a) GDSC schweinfurthin A activity versus gene expression-based EMT index value. Red points indicated cell lines with intermediate ‘epithelial-mesenchymal’ status, while remaining points on the left and right are classified as mesenchymal and epithelial, respectively. (b) Activity of schweinfurthin A vs. activity of 5-methylschweinfurthin G in a subset of GDSC cell lines. (c) Bar plot of Pearson’s correlations between GDSC drug activities and EMT index.

**Supplementary Figure S5:**
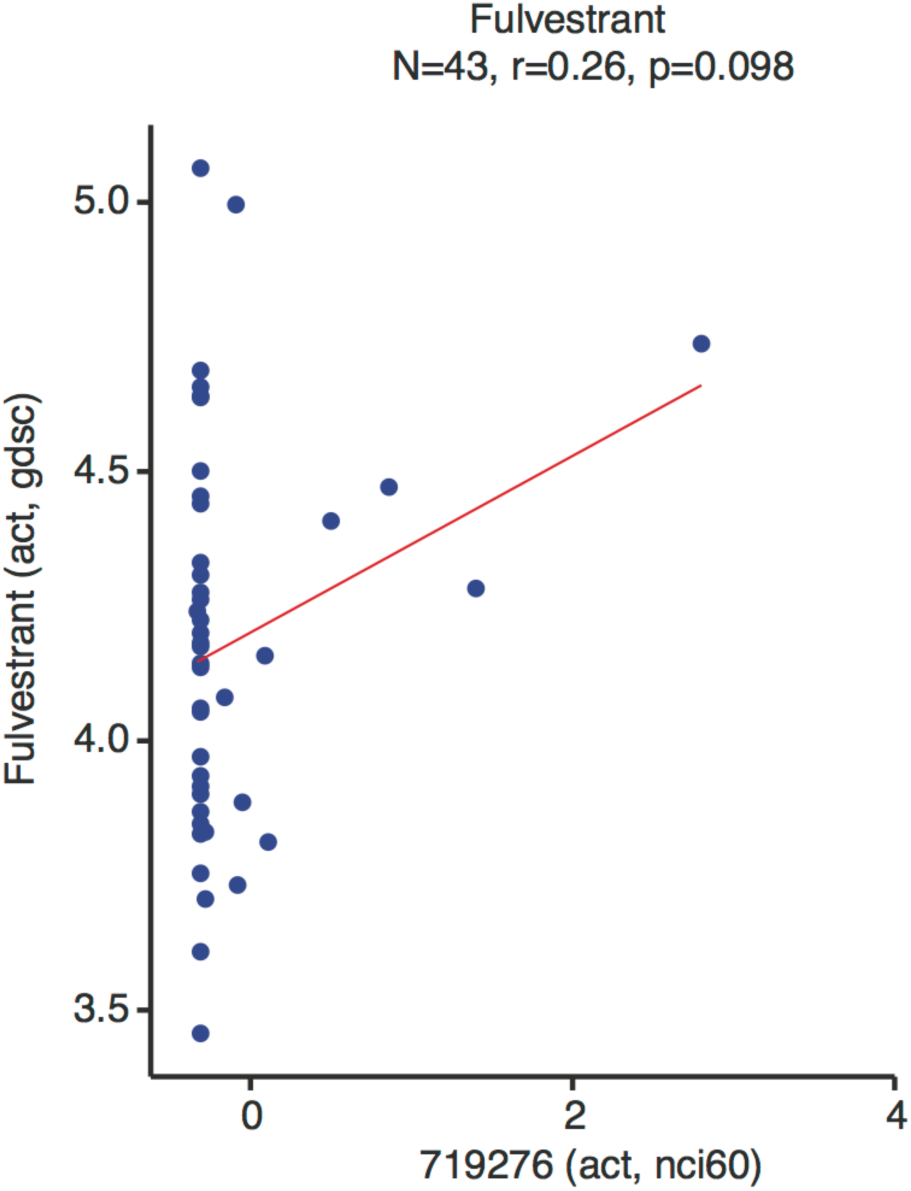
GDSC versus NCI-60 drug activity for fulvestrant, indicating inappropriate drug concentration range in NCI-60 activity assay.

**Supplementary Figure S6:**
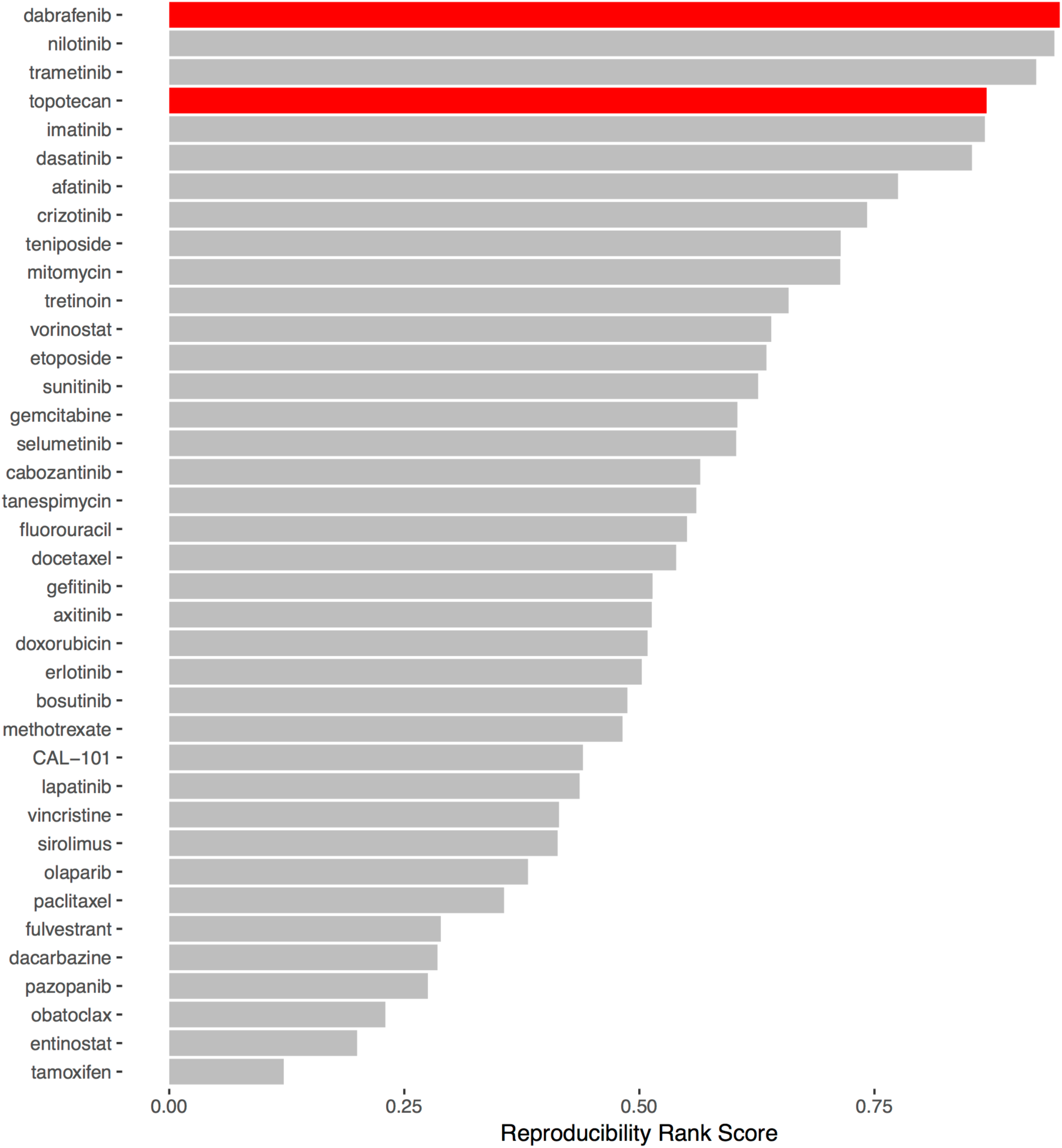
Overall drug activity data reproducibility rankings for 38 compounds tested in the NCI-60, GDSC, and CTRP, integrating pairwise activity correlations between the sources.

